# Effect of sleep deprivation on fractal and oscillatory spectral measures of the sleep EEG: a window on basic regulatory processes

**DOI:** 10.1101/2024.08.29.610292

**Authors:** Csenge G. Horváth, Róbert Bódizs

## Abstract

Sleep is vital for sustaining life; therefore, reliable measurement of its regulatory processes is of significant importance in research and medicine. Here we examine the effect of extended wakefulness on the putative indicators of fundamental sleep regulatory processes (spectral slope and spindle frequency) proposed by the Fractal and Oscillatory Adjustment model of sleep regulation by involving a healthy young adult sample in a 35-hour long sleep deprivation protocol. Wearable headband EEG-derived results revealed that NREM sleep electroencephalogram (EEG) spectral slope estimated in the 2–48 Hz range is an accurate indicator of the predicted changes in sleep depth induced by sleep deprivation (steepened slopes in recovery sleep) or by the overnight dissipation of sleep pressure (flattening slopes during successive sleep cycles). While the baseline overnight dynamics of the center frequency of the sleep spindle oscillations followed a U-shaped curve, and the timing of its minimum (the presumed phase indicator) correlated with questionnaire-based chronotype metrics as predicted, a different picture emerged during recovery sleep. Advanced recovery sleep advanced the timing of the minima of the oscillatory spindle frequency, reduced considerably its relationship with chronotype, but retained partially its U-shaped overnight evolution. Overall, our study supports the use of the spectral slope of the sleep EEG as a homeostatic marker of wake-sleep regulation, in addition, encourages further research on the EEG-derived measure of the circadian rhythm, primarily focusing on its interaction with the homeostatic process.

## 1. Introduction

In the past few years, the characterization and appropriate parametrization of the EEG Fourier power spectrum has gained increasing interest, as it become clear that besides the undeniable importance of neural oscillations, the aperiodic brain activity also carries substantial information on neural functioning. The same holds true for the field of sleep research, where nowadays -after the proper distinction of the periodic and aperiodic part of the spectrum-, analyses of the aperiodic component came to the forefront.

The so-called spectral slope (or spectral exponent) describes the linear relationship between the logarithm of power and the logarithm of frequency which is due to the power-law distribution of the EEG spectrum (Bódizs et al., 2021; Feinberg et al., 1984; Pritchard, 1992). In a recent paper of ours, we thoroughly reviewed the literature and came to the conclusion that results regarding the spatio-temporal and age-related variation of the sleep EEG spectrum are highly consistent with the assumption of the pivotal role of aperiodic activity in wake-sleep regulation (Bódizs et al., 2024). The EEG metrics of aperiodic activity were found to be a suitable indicators of consciousness and arousal (Colombo et al., 2019; Lendner et al., 2020; Waschke et al., 2021; Zhang et al., 2023), as well as reliable measures in distinguishing between different brain states like resting and active wakefulness or between stages of sleep (Höhn et al., 2024; Miskovic et al., 2019; Schneider et al., 2022). Additionally, we found that the temporal evolution of the spectral slope throughout the night depends on the preliminary amount of sleep, that is slope flattens as sleep progresses (G. Horváth et al., 2022), based on which it is a putative indicator of sleep-wake history. Hence, results of the literature support the assumption that EEG spectral slope could be a reliable marker of sleep homeostasis. To put it simply, sleep homeostasis refers somehow to the constancy of the sleep-wake dynamics. When one state overloads, the other has to recover so that the system can be stable again.

However, to maintain a balanced functioning within the context of ecologically definitive day-night cycles, the timing of sleep and wake is also an important factor. Former research indicate that sleep spindle frequency can convey information on this timing when EEG data is considered. Spindle frequency was found to covariate with reliable circadian indicators, like core body temperature and melatonin (Knoblauch et al., 2005; Wei et al., 1999), furthermore, the timing of the nadir of the spindle frequency (NSSF) associated with actigraphy-derived circadian phase index (G. Horváth & Bódizs, 2024). Time-of-day dependence of spindle frequency activity was proved directly with specific study designs targeting the separated analysis of circadian phase and sleep-wake history (Aeschbach et al., 1997; Dijk, 1999). These studies found larger spindle activity at lower spindle frequencies during the night-time and this activity dominance transferred to higher frequencies towards daytime. Further indirect proofs of time-of-day-dependency are the results showing spindle frequency is higher during daytime as compared to night-time sleep (Bódizs et al., 2022; Rosinvil et al., 2015), and follows a U-shape curve during the night with the minimum around the middle of the sleep period (Bódizs et al., 2022; G. Horváth et al., 2022; G. Horváth & Bódizs, 2024).

The ground-breaking framework which considers the interaction of timing and sleep homeostasis as the foundations of sleep regulatory mechanisms is the two-process model of sleep regulation proposed by Alexander Borbély (Borbély, 1982). However, the original model relies only on the oscillatory activity of the brain, and no EEG indicator for the circadian process (that is for the timing) was suggested. In our recent work, we recommended a new, so-called Fractal and Oscillatory Adjustment Model by revising the two-process model (Bódizs et al., 2024). In this work, we took into account the discovery that periodic and aperiodic components are two types of coexisting brain activity with equally important information content. Besides this model proposition, our aim was to support the assumptions that sleep homeostasis is reliably reflected by the spectral slope while circadian regulatory process can be indexed by the oscillatory frequency in the spindle band.

To validate the above assumptions, we implemented the recordings and analyses of baseline- and recovery sleep (BS and RS, respectively) of healthy young adults with the use of a 35-hour long home sleep deprivation protocol relying on wearable headband EEG devices. To our best knowledge this is the first report in which spectral slope and oscillatory spindle frequency were examined by experimental challenge of the homeostatic process (35-hours of sleep deprivation) and with a slight rescheduling of sleep with the aim of unravelling the circadian clock (RS period advanced by ∼4.5 hours). We parametrized the NREM EEG power spectra for every sleep cycle (NREM-REM periods), furthermore, we defined spindle frequency as the center frequency of the most prominent peak in the 9-16 Hz frequency range as indicated by the parametrization procedure.

Sleep recordings were analysed in detail with multiple approaches. Besides the differences of BS and RS, the overnight dynamics of spectral slopes and peak center frequencies in the spindle frequency range (CF) were tested. These analyses were performed by involving the whole sample, which approach limited the analysable number of sleep cycles to the first 4 of the nights. However, overnight dynamics were further analysed by grouping participants according to their total number of sleep cycles, in which case the group sizes were limited. Furthermore, regarding the comparison of the two sleep conditions, specific sleep cycles (first, middle, last), in addition, maximal and minimal values of the parameters and their occurrences (cycle number, or timing) were also analysed.

### Based on the above, we tested the following hypotheses

A. Comparison of the two sleep condition will show direct evidence for the involvement of the two parameters in sleep regulatory processes. More precisely, we hypothesize that:
  hyp 1. Slope will be flatter in BS than in RS in the first 4 cycles of sleep (experimental challenge of the sleep homeostat results in increased sleep EEG spectral slope steepness)
  hyp 2. Spectral EEG slope values of the BS and RS conditions will differ to a greater extent in the first cycles, as compared to the last cycles of sleep due to the increased sleep pressure caused by the deprivation at the beginning of the night, as well as because of the concept of homeostasis assumes that a near-equilibrium state should return at the end of recovery sleep.
  hyp 3. Steepest slope will occur at the beginning of the night and this slope will be flatter in BS than in RS. However, there will be no difference regarding the minimal values (same explanation as for hyp 2) which are hypothesized to occur at the end of the night in both conditions.
  hyp 4. The development of CF in the first 4 cycles of sleep in BS and RS will be different because of the different times of day in which they fall.
  hyp 5. CF values will not differ in the last and first cycles of the two nights
  hyp 6. CF minima will appear around the same time in both nights, that is NSSF will not differ between conditions as it depends more on time-of-day. Furthermore, NSSF in BS (when the sleep schedule is regular) will occur around the middle of the sleep period.
B. Throughout its overnight dynamics, slope and CF will reflect homeostatic and circadian processes respectively, as:
  hyp 7. Slope will flatten during the night in both sleep condition (BS and RS) in parallel with the decrease of sleep pressure in the first 4 cycles of sleep
  hyp 8. The flattening trend of slope values will persist even if the whole sleep period is considered
  hyp 9. CF will evolve according to a U-shaped curve (decelerate in 2^nd^ or 3^rd^ cycle) in the first 4 cycles of sleep in BS, but not in RS. We hypothesized this latter is caused by the advanced sleep schedule (sleeping period starts approximately 4.5 hours earlier than the habitual bedtime of the participants), as well as because of the heightened number of sleep cycles in RS (the 4^th^ cycle should fall around the middle of the night).
  hyp 10. There will be a deceleration (U-shape-like evolution) of CF at the middle of the night, in both BS and RS, when the whole sleep period is considered.

## 2. Methods

### 2.1. Sample

In this experimental study N = 46 healthy young adults participated in a 7-day long protocol involving actigraphy, headband-wearable recorded sleep EEG, and a 35 hour-long sleep deprivation followed by ad libitum recovery sleep. Due to poor quality EEG recordings or complete data loss the final sample consisted of N = 38 subjects (age range: 18–39 years, mean age= 24.9, 19 females).

Participants were enrolled by a combination of convenience and snowball sampling procedures involving personal contacts and social media calls. All subjects were free of psychiatric or neurological disorders based on self-reports. In addition, the exclusion criteria of the study included the Hungarian version of Pittsburgh Sleep Quality Index (Takács et al., 2016) score over 5, Beck Depression Inventory (Beck et al., 1961) score over 12 (moderate and severe depression symptoms (Rózsa et al., 2001), alarm clock usage on free days, extreme circadian preference (MCTQ chronotype scores outside of the +/-2 SD of reported values in young Hungarian subjects according to Haraszti et al. (2014) and shift work, as well as reported acute and/or chronic medical diagnoses or ongoing pharmacological treatments.

### 2.2. Study protocol

After the questionnaire-based testing of inclusion/exclusion criteria, enrolled participants had to wear a GENEActiv triaxial accelerometer (Activinsights, Cambridge, United Kingdom) for 7 days on their non-dominant wrist. No instructions were given for the first 4 days except not to take off the actigraphy device for any activity. On the 5th day, participants had to come to the laboratory to learn how to use the mobile EEG device and to perform computerized cognitive tasks. On this same evening (5th night of the experiment), subjects slept with the wearable EEG headband in their own home. During this BS EEG measurement, the bedtime was freely chosen by the participants thus, they could go to bed according to their own preferences. The use of an alarm clock was prohibited in both the mornings of headband-EEG-recorded sleep (BS and RS). After waking up from BS, the sleep deprivation part of the study began. Participants had to stay awake for 35 hours during which alcohol and any kind of stimulant consumption was prohibited with the exception of caffeine. Participants were allowed to consume as much coffee as they typically would in an average 24-hour day; however, caffeine intake was restricted to no more than the equivalent of 3 espresso shots during the entire sleep deprivation period. As it was an at-home examination, participants could freely choose their activities throughout the 35 hours of wakefulness, however, they had to have continuous contact with the experimenter (at least one report on their well-being and actual activity per hour via mobile phone messages) and complete questionnaires (sleepiness scales) at specified time points (in the 0., 12., 24., and 35. hour of the sleep deprivation). Moreover, in the 24^th^ and 34^th^ hour of sleep deprivation, experimenters went to the participants’ homes to record the cognitive tasks and check their well-being and their adherence to the study protocol in person. Overall, compliance with the study protocol was checked by verifying whether all hourly text messages were sent, and the questionnaires were completed at every given time points by the participants. Finally, compliance was confirmed through offline post-participation evaluation of actigraphy data if there were no indications of sleep during the 35-hour period. At the end of the sleep deprivation, the experimenter helped the subjects to put on the EEG headband and instructed them to turn off the lights and go to bed. This was the recovery night sleep EEG measurement. After waking up, participants’ last task was to fill out the sleepiness scales one more time, then the experiment ended. In the present study some of the questionnaires (MCTQ, sleepiness scales) and the sleep EEG data were analysed.

National Public Health Centre Institutional Committee of Science and Research Ethics approved the research protocols, and the experiment was implemented in accordance with the Declaration of Helsinki. Every participant signed an informed consent about their attendance in the study.

### 2.3. Sleep EEG

Electroencephalography of sleep was recorded by Hypnodyne corp. Zmax EEG headband with sampling rate of 256 Hz at derivations F7-Fpz and F8-Fpz re-referenced to their common average. Manual sleep scoring and artefact removal was performed in 20 and 4 second epochs, respectively. Sleep recordings was divided by sleep cycles (successive NREM+REM episodes) and only NREM sleep was analysed in each cycle. Determination of sleep cycles was based on the modified Feinberg and Floyd criteria (Feinberg & Floyd, 1979) proposed by Jenni & Carskadon (Jenni & Carskadon, 2004) with the consideration of skipped REMs in the first cycle of sleep.

### 2.4. Parametrization of the sleep EEG spectrum

The fitting oscillations and 1-over-f (FOOOF) method, developed by Donoghue et al. (2020), aims to parametrize the power spectrum into aperiodic and periodic components. The method can be described briefly with the following steps. First, it estimates and removes an approximated aperiodic component from the spectrum, thus creating a flattened version of the spectrum that highlights the spectral peaks. Then, Gaussians are iteratively fitted and subtracted to isolate the periodic components. Periodic components are removed from the spectrum, and the aperiodic part is refitted, then the combined version of the periodic and refitted aperiodic component can be used for further analyses.

To avoid over- or under fitting, the settings recommended by Schneider et al. (2022) was used in the present study. Fitting range was set to the 2–48 Hz frequency interval, bandwidth of accepted peaks was 0.7–3 Hz, and the peak threshold was set to 1.

### 2.4. Questionnaires

#### 2.4.1. Munich Chronotype Questionnaire

MCTQ collects information on usual bed- and wake times, sleep latency and inertia, separately for working and free days. From these time points, the middle of the sleep periods is calculated. Finally, chronotype is given as the oversleeping adjusted sleep midpoint on the free days (MSFsc) (Roenneberg et al., 2003). In most cases, this variable was used as a continuous predictor. However, groups of the different chronotypes (early-, intermediate-, and evening types) were also calculated. A person was classified either early or evening types if their values fell outside the ± 1 SD range (but within ±2 SD) and as intermediate if their MSFsc fell inside the ± 1 SD range of the mean of young Hungarian subjects reported by Haraszti et al (2014).

#### 2.4.2. Sleepiness scales

Sleepiness was measured by two scales at the above-mentioned time-points. Firstly, by using a simple likert-scale from 1 to 10 which was reported more sensitive to daytime dysfunction than other subjective rating scales (Riegel et al., 2013): participants were asked to rate their sleepiness level where 1 meant ‘not sleepy at all’ while 10 referred to ‘very sleepy’. Secondly, with the widely-used Stanford Sleepiness Scale (Hoddes et al., 1973). The instrument consists of 7 phrases describing sleepiness, and participants are asked to choose the one which best indicates their current sleepiness level.

### 2.5. Statistics

All statistical analyses were carried out using TIBCO Statistica software (TIBCO Software Inc., 2020). Data examination was performed in multiple settings.

We were interested in the overnight dynamics of specific spectral EEG parameters during the two nights (BS and RS) separately, as well as in the comparison of BS vs. RS. Both research questions were analysed in two ways. Firstly, in a simplified manner where we strived for keeping the sample size as high as possible, thus, the number of analysable sleep cycles were limited (the first 4 cycles were examined). Furthermore, we grouped participants according to their number of sleep cycles, and evaluated the overnight dynamics in these groups separately. In this case the sample sizes were low in the groups, but we could assess the overnight dynamics for the total night of each participant who had complete recordings. As the cycle effect was hypothesized to be substantial, we considered a group analysable if its sample size was at least N = 8, consequently the BS groups with 3 and 7, furthermore, RS groups with 6 and 10 sleep cycles had to be excluded as these consisted of a maximum of 3 participants. Finally, specific sleep cycles (at the beginning, the middle, and the end of the night sleep periods), as well as the sleep cycle numbers corresponding to the minimum and maximum values of the parameters were compared between the two nights.

Differences between variables and conditions were tested with the appropriate parametric statistical tests on normally distributed data: dependent sample t-test for two variable/condition comparison, Repeated Measures ANOVA where more than two variables/time-points were involved, General Linear models in case of multiple factors included. If the sample size was less than 10 (cycle group-wise analyses) or one of the variables included in the analysis had non-Gaussian distribution, non-parametric statistical version of the above-mentioned tests were used: Wilcoxon Matched pairs test, Friedman ANOVA. In the descriptive statistics mean (m) and standard deviation (SD) are reported for data with normal distribution, and median (Mdn) with interquartile range (IQR=[lower border, upper border]) for non-normally distributed data. Test of normality was performed by Shapiro-Wilk W test for all variables. According to its results, the following variables significantly deviated from a normal distribution: the number of sleep cycles in both BS and RS, the sleep cycle in which participants had their CF minimum in BS, slope in BS cycle 4 and RS cycle 7, both sleepiness scales at all time-points, the cycles which contain the BS and RS slope value minimum and maximum.

The threshold of significance was defined as p < 0.05 in all analyses, in addition, Bonferroni correction was applied with the appropriate significance level, where multiple comparisons were conducted.

## 3. Results

Out of the recordings of the 38 participants 3 in the baseline and 7 in the recovery nights were incomplete due to device failure (e.g. poor quality data, electrode contact problems, unexpected shutdown of the device). These recordings were not involved in the analyses which assume the total length of the nights (e.g. total night dynamics, minimum/maximum values during the night, circadian analyses), but included in the cycle-based (sample mean) comparisons between BS and RS.

Participants stayed awake Mdn=35.79 hours (∼35 h. and 47 min; min-max range: 34.9–37.3). This was estimated from the scored recordings of BS and RS (length of the period between BS last sleep episode and RS sleep initiation for participants whose BS recording was complete). The average sleep duration: m_BS_=7.9 (min-max: 5.8–10.7) hours, m_RS_=12.2 (min-max:9.7–14.74) hours, as well as the sample median of sleep cycles Mdn_BS_=5 (min-max:3-7) and Mdn_RS_=8 (min-max: 6–10) were also calculated for complete recordings.

The sample mean of MSFsc (MCTQ-based chronotype metric) was at 04:34 (SD=1:16 hh:mm). In total, 7 early-, 15 intermediate-, and 16 evening type (5, 6, 8 females respectively) subjects’ data were analysed. Participants slept according to their circadian timing in BS as BS sleep start (derived from the EEG recordings) strongly positively correlated with the MCTQ-based chronotype results (r=0.61, p<0.0001). However, the correlation between the questionnaire and RS sleep starts was weaker (r=0.3, p=0.07; Fig. 1/A). Sleepiness level increased the most from 12^th^ to the 24^th^ hour of the sleep deprivation period according to both scales and reduced again after waking up from RS (Figure 1/B). The difference in sleepiness levels between successive time points is significant from the 12^th^ to the 24^th^ hour (t_likert_(37)=-9.78, t_SSS_(37)=-9.82, p<0.0001) and from the 35^th^ hour to RS wake (t_likert_(37)=6.81, t_SSS_(37)=6.2, p<0.0001).

**Figure 1.**
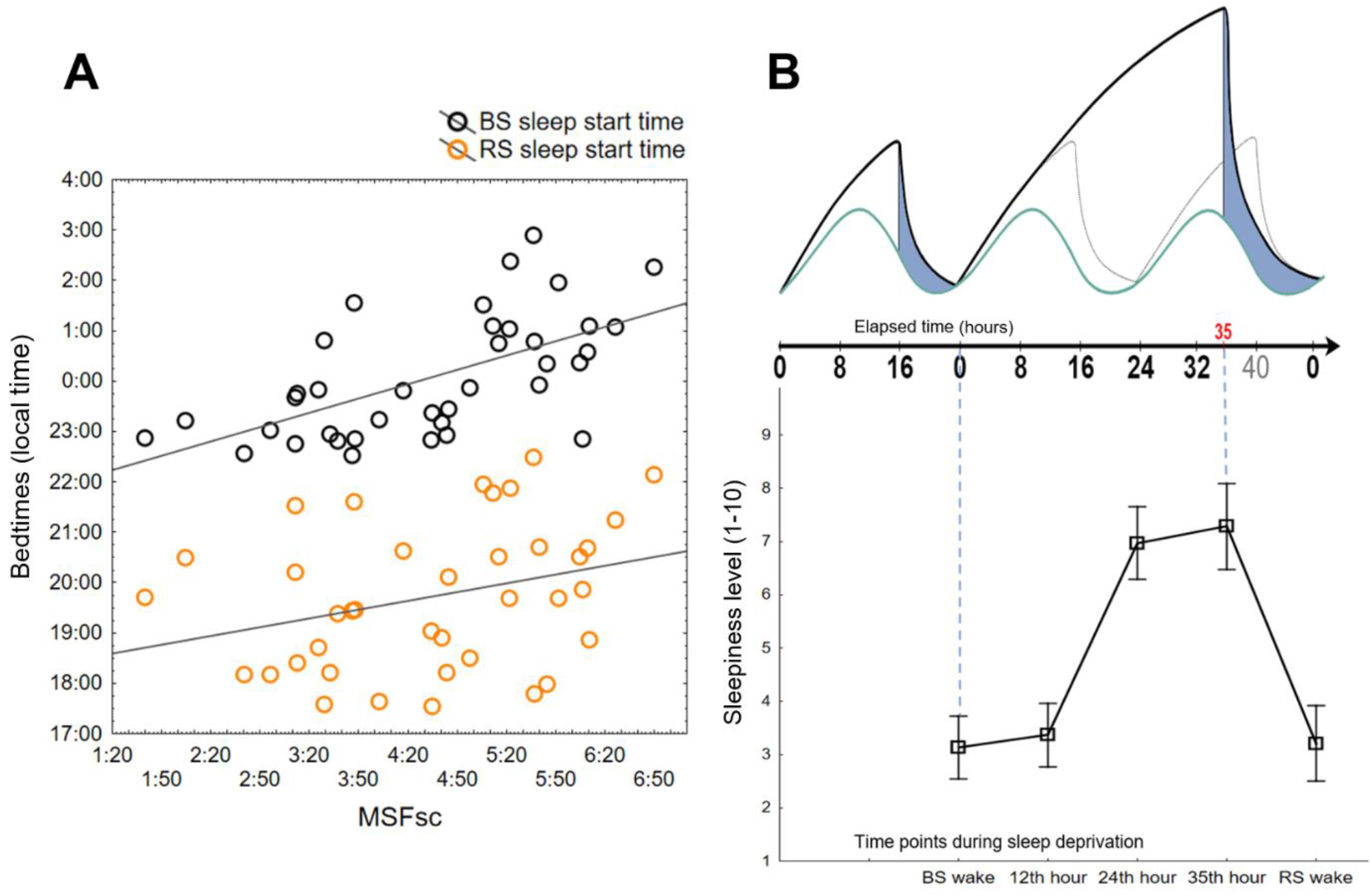
Subjective chronotype and sleepiness along with the expected sleep pressure and circadian time according to the two-process model of sleep regulation. A) Correlation between the MCTQ chronotype indicator (MSFsc) and actual bedtimes of BS and RS. B) Schematic representation of the expected behaviour of the two sleep regulatory processes (top) and the actual development of sleepiness according to the likert scale (bottom) during the experiment. On the top, blue area indicates the approximate time duration of the sleep periods, while the black and light-grey lines demonstrate the homeostatic sleep pressure due to the intervention, and under normal circumstances without sleep deprivation, respectively (note the heightened sleep pressure due to the extended wakefulness). On the bottom, the graph displays the sample means with 95% confidence intervals of sleepiness levels at different time points during the wakefulness.

### 3.1. EEG spectral slope

Effect of sleep deprivation was tested on the whole sample with General linear model using cycle and sleep condition (RS vs. BS) as within subject factors. Spectral slope was significantly steeper after sleep deprivation (hyp 1; F=65.94, p<0.0001) and flattened in the first 4 cycles of sleep (hyp 7; F=98.25, p<0.0001; Table 1), furthermore the condition × cycle interaction was also significant (F=2.95, p=0.037, Fig. 2.). Unequal N HSD posthoc test was conducted to check whether successive sleep cycles were differed from each other in spectral slope values. Sample-level slope difference (Fig. 2.) was significant between C2 vs. C3 (BS & RS: p<0.001,) and C3 vs. C4 (BS: p=0.02, RS: p<0.001) in both conditions, but not between C1 vs. C2 (BS: p=1.0, RS: p=0.83).

**Table 1.** Sample means and standard deviation of slope values in the first four sleep cycles.

| cycle | baseline |  |  | recovery |  |  |
| --- | --- | --- | --- | --- | --- | --- |
|  | N | Mean | SD | N | Mean | SD |
| 1 | 36 | 2.402602 | 0.126357 | 38 | 2.625733 | 0.214957 |
| 2 | 36 | 2.388576 | 0.170833 | 36 | 2.615997 | 0.199933 |
| 3 | 37 | 2.218177 | 0.205926 | 34 | 2.410661 | 0.189354 |
| 4 | 36 | 2.100263 | 0.179556 | 33 | 2.247819 | 0.129501 |

**Figure 2.**
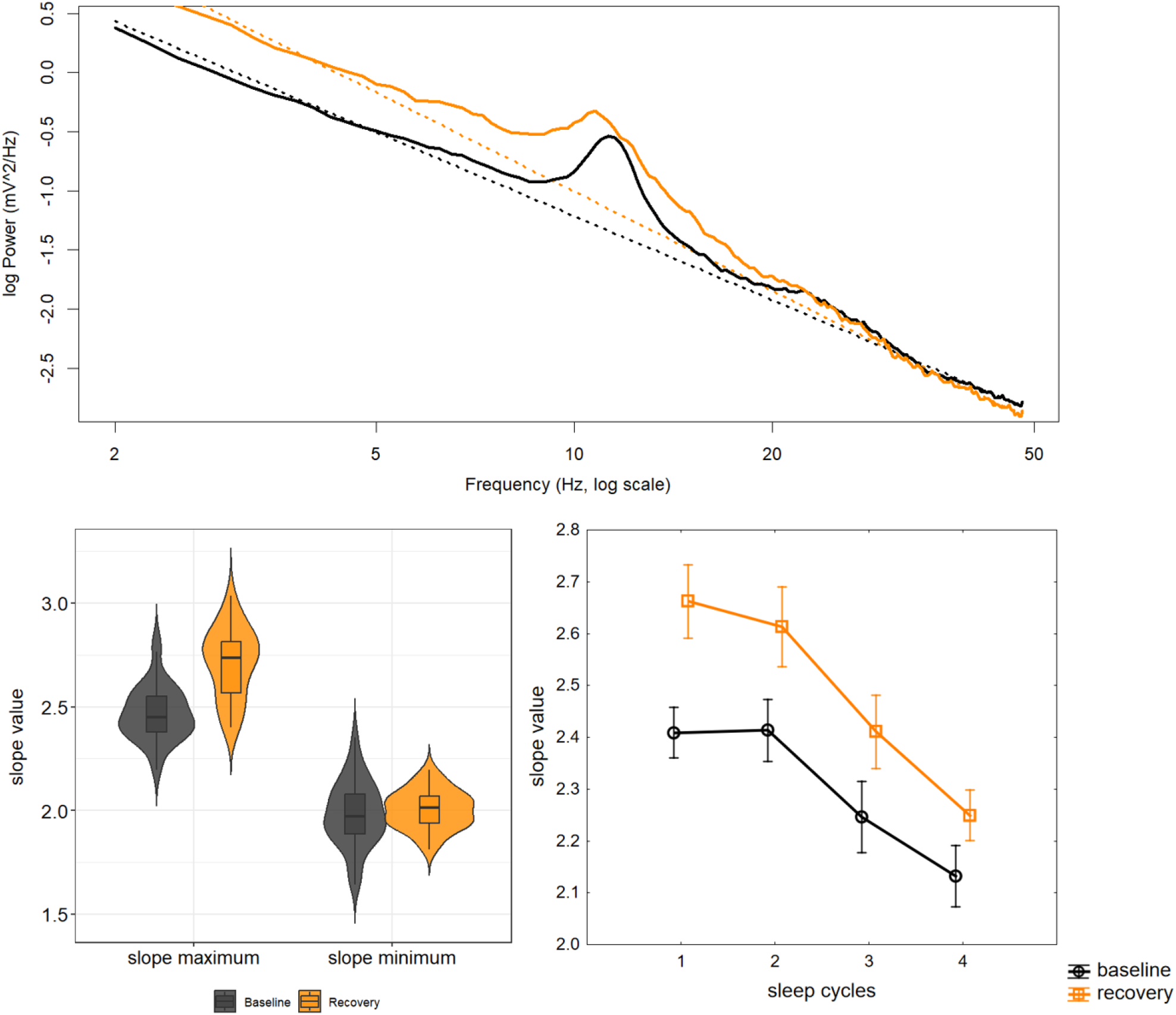
Spectral slopes of the NREM sleep EEG during the first four sleep cycles of the baseline and the recovery nights. Upper panel shows the spectrum of the first cycle of sleep for BS (black) and RS (orange), as well as the fitted aperiodic components with dashed lines in a 24-year-old male participant. Violin plots (left side of the lower panel) depict the distribution of the maximal and minimal slope values, whereas inner boxplots show the difference between sleep conditions: while maximum slope values got larger due to the sleep deprivation, slope minima returned to approximately the same level in the two conditions. Right panel shows the sample means and 95% confidence interval of slope values in the first 4 cycle of sleep in BS and RS. Slope is steeper in RS than in BS at the beginning of the sleep.

**Figure 3.**
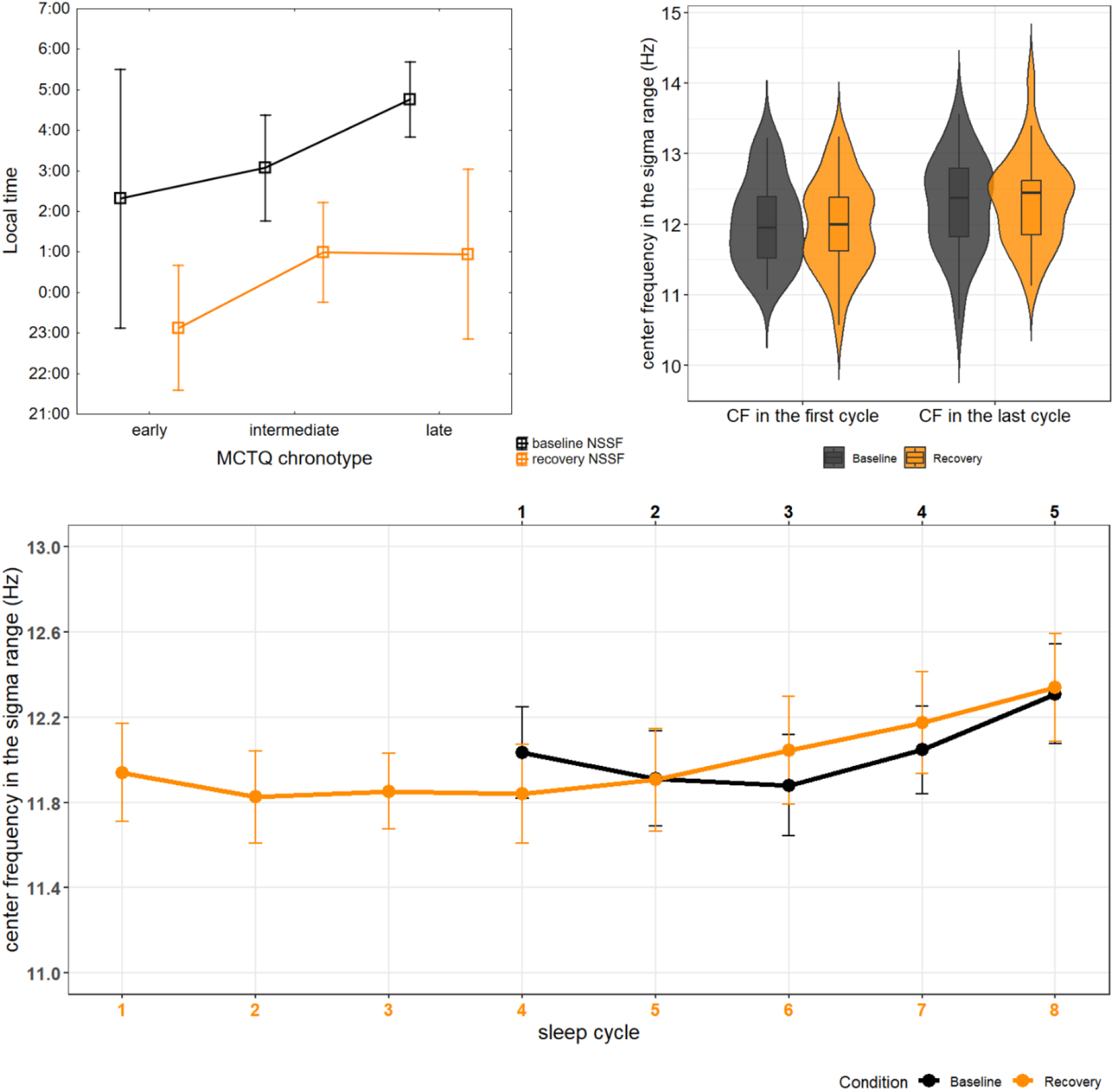
Difference in the CF minima (NSSF) in RS and BS between MCTQ chronotype groups, and in CF values between sleep conditions. Subjectively measured chronotype is well reflected in the time of BS NSSF, but not in RS NSSF, additionally, NSSF times differ between sleep conditions (upper left panel). Distribution, and medians of CF in the first and last cycles of the sleep episodes are not different (violin/boxplot on the right). Lower panel represents the sample means and 95% confidence intervals of CF in successive sleep cycles in both conditions and displays the shifted sleep period which was advanced by approximately 3 sleep cycles.

In order to further analyse the overnight dynamics of the whole sleep period and the differences of BS and RS, group-level and cycle-specific analyses were conducted.

Cycle count-based group analyses showed that the cycle effect remained significant even if the whole sleep period was considered (hyp 8; Table 2; Fig.4.). To check whether successive sleep cycles were differed from each other Wilcoxon matched-pairs test was carried out, followed by Bonferroni correction for multiply comparisons resulting in a significance levels set at p<0.0125, p<0.01, p<0.008, p<0.007, p<0.00625, p<0.00555, and p<0.005 for 4, 5, 6, 7, 8, 9, and 10 comparisons, respectively. After the correction, no significant difference in BS spectral slopes between successive cycles (e.g. cycle 1 vs. cycle 2, cycle 2 vs. cycle 3, …, cycle 6 vs. cycle 7) was revealed (in none of the cycle count based groups). However, RS spectral slope differences remained significant in the 9-cycle group between C2 and C3 (Z=2.8, p=0.0051).

**Table 2.** Overnight cycle effect of slope and CF values in the separate cycle count-based groups. Friedman ANOVA revealed significant or marginally significant cycle effect in all groups except with regards CF in the BSC4 group.

| Groups defined by maximum cycle count | Sample size | Slope |  | CF |  |
| --- | --- | --- | --- | --- | --- |
| | | $\chi^2$ | p | $\chi^2$ | p |
| BSC4 | 7 (6) | 7.3 | 0.06 | 1.8 | 0.61 |
| BSC5 | 12 (10) | 33.33 | <0.0001 | 13.5 | 0.009 |
| BSC6 | 11 | 37.13 | <0.0001 | 23.94 | 0.0002 |
| ReSC7 | 8 (7) | 41.3 | <0.0001 | 11.51 | 0.07 |
| ReSC8 | 8 (7) | 47.54 | <0.0001 | 22.86 | 0.002 |
| ReSC9 | 10 | 60.27 | <0.0001 | 34.24 | <0.0001 |

Furthermore, we compared the parameter in the first and last cycle of the sleep periods (BS C1 vs. RS C1, BS Clast vs. RS Clast) as we hypothesized that the main difference will be at the beginning of the night (hyp 2). Indeed, in the first cycle of BS the slope was flatter than in RS (hyp 3; BS: m=2.4, SD=0.1; RS: m=2.63, SD=0.2, t(35)=-6.6, p<0.0001), whereas, slope values of the last cycles were not significantly different in the two conditions (BS: m=2.06, SD=0.15; RS: m=2.08, SD=0.14; t(29)=-0.5, p=0.62). As a next step, we defined the maximum and minimum slope values along with their corresponding sleep cycle numbers to support the assumption that the steepest and flattest values are at the beginning and at the end of the sleep period, respectively. Besides, we compared them between sleep conditions in order to test the hypothesis stating that the BS vs RS difference in slope maxima will be larger than the divergence in the minima (hyp 2). In BS, slope was the steepest in the first cycle of sleep for 16, in the 2^nd^ cycle for 17, and in the 3^rd^ for 2 subjects; in RS, 18 subjects had their steepest slope in the 1^st^ sleep cycle, and 13 in the 2^nd^ (remaining participants did not have complete recordings). Slope maximum was significantly steeper in RS (t(29)=-9.5, p<0.0001), furthermore, BS and RS did not differ significantly in terms of which sleep cycle contained the steepest slope value (Z=0.2, p=0.23, BS: Mdn=2 (2nd cycle), IQR=[1,2]; RS: Mdn=1(1st cycle), IQR=[1,2]). Minimum slope values did not differ between BS and RS (t(29)=-1.2, p=0.24), moreover, the cycle number of the last cycles significantly differed from the cycle number in which the minimum values (flattest slopes) occurred, that is flattest slope did not always fall in the last sleep cycle of the night (**BS**: Z=4.2, p<0.0001, *num. of lastC*: Mdn=5, IQR=[5, 6], *num. of MinC*: Mdn=4.5, IQR=[3, 5]; **RS**: Z=3.9, p<0.0001, *num. of lastC*: Mdn=8, IQR=[7, 9], *num. of MinC*: Mdn=7, IQR=[6, 8].

### 3.2. Spindle frequency

General Linear Model with condition (BS/RS) and cycle as within subject factors, showed that when the first 4 cycle of the whole sample were considered (hyp 4), only cycle affected significantly CF (BS/RS: F(1,27)=3.13, p=0.1, cycle: F(3,81)=3.27, p=0.025). The interaction was not significant.

Similar to the approach used for spectral slope analyses, further details on the overnight dynamics of the total night in the separate cycle count-based groups, as well as CF similarities/differences of the two sleep conditions were analysed.

Cycle effect in CF was significant in all groups except the 4-cycle group in BS and 7-cycle group in RS (Fig. 4., see descriptive statistics in Table 3). However, as we hypothesized that CF will decelerate around the middle of the night sleep period, we conducted sample-level comparisons between the middle- and first-, as well as between the middle- and last sleep cycles’ CF values using dependent sample t-test with Bonferroni correction for multiple comparisons, resulting in a significance level set at p<0.025 for the 2 comparisons. Where the maximum cycle count was an even number, average of the two middle cycles was considered as the middle of sleep. In the middle of the sleep period CF was significantly slower than in the last sleep cycle in both conditions (**BS**: *C*_*middle*_ [m=11.85, SD=0.7] vs. *C*_*last*_ [m=12.25, SD=0.7]: t(34)=-3.8, p<0.001; **RS**: *C*_*middle*_ [m=11.8, SD=0.66] vs. *C*_*last*_ [m=12.31, SD=0.6]: t(30)=-5.3, p<0.0001), however, sleep middle CFs did not differ significantly from CFs of the first sleep cycles (**BS**: *C*_*first*_ [m=12.03, SD=0.6] vs. *C*_*middle*_ : t(32)=1.86, p=0.07; **RS**: *C*_*first*_ [m=11.93, SD=0.7] vs. *C*_*middle*_ : t(28)=1.04, p=0.31). After that, we compared last and first CF values in the sample to check whether the beginning and the end of sleep is similar regarding spindle CF. Last cycle CFs were significantly faster than first cycle CFs in both condition (**BS**: t(33)=-3.8, p<0.001; **RS**: t(28)=-4.2, p<0.001).

**Table 3.** Sample sizes, medians and interquartile ranges of CF values in the different cycle count-based groups.

| Cycle | N | Mdn | IQR:25% | IQR:75% | N | Mdn | IQR:25% | IQR:75% | N | Mdn | IQR:25% | IQR:75% |
| --- | --- | --- | --- | --- | --- | --- | --- | --- | --- | --- | --- | --- |
|  | **BSC4** |  |  |  | **BSC5** |  |  |  | **BSC6** |  |  |  |
| 1 | 7 | 12.20 | 11.26 | 12.47 | 11 | 12.01 | 11.74 | 12.39 | 11 | 11.76 | 11.57 | 12.31 |
| 2 | 6 | 11.92 | 10.98 | 12.28 | 10 | 11.93 | 11.68 | 12.32 | 11 | 11.78 | 11.33 | 12.25 |
| 3 | 6 | 11.99 | 11.15 | 12.24 | 12 | 11.95 | 11.38 | 12.35 | 11 | 11.74 | 11.41 | 12.24 |
| 4 | 7 | 12.29 | 10.96 | 12.53 | 12 | 12.27 | 11.79 | 12.61 | 11 | 11.84 | 11.53 | 12.29 |
| 5 |  |  |  |  | 12 | 12.47 | 11.89 | 12.87 | 11 | 11.94 | 11.59 | 12.52 |
| 6 |  |  |  |  |  |  |  |  | 11 | 12.11 | 11.74 | 12.67 |
| 7 |  |  |  |  |  |  |  |  |  |  |  |  |
|  | **RSC7** |  |  |  | **RSC8** |  |  |  | **RSC9** |  |  |  |
| 1 | 8 | 11.69 | 11.33 | 12.08 | 7 | 12.22 | 11.64 | 12.46 | 10 | 11.95 | 11.62 | 12.38 |
| 2 | 7 | 11.53 | 11.37 | 12.19 | 7 | 11.93 | 11.89 | 12.22 | 10 | 11.60 | 11.34 | 12.29 |
| 3 | 7 | 11.50 | 11.29 | 12.03 | 7 | 11.78 | 11.60 | 12.17 | 10 | 11.88 | 11.62 | 12.14 |
| 4 | 8 | 11.89 | 11.06 | 12.34 | 8 | 11.72 | 11.25 | 12.09 | 10 | 12.06 | 11.49 | 12.11 |
| 5 | 8 | 11.65 | 11.10 | 12.18 | 8 | 11.87 | 11.65 | 12.62 | 10 | 11.85 | 11.23 | 12.22 |
| 6 | 8 | 11.92 | 11.28 | 12.38 | 8 | 12.08 | 11.80 | 12.64 | 10 | 12.05 | 11.11 | 12.73 |
| 7 | 8 | 12.06 | 11.57 | 12.52 | 8 | 12.44 | 11.88 | 12.99 | 10 | 12.00 | 11.59 | 12.38 |
| 8 |  |  |  |  | 8 | 12.57 | 11.92 | 12.82 | 10 | 12.04 | 12.01 | 12.43 |
| 9 |  |  |  |  |  |  |  |  | 10 | 12.31 | 11.87 | 12.60 |

**Figure 4.**
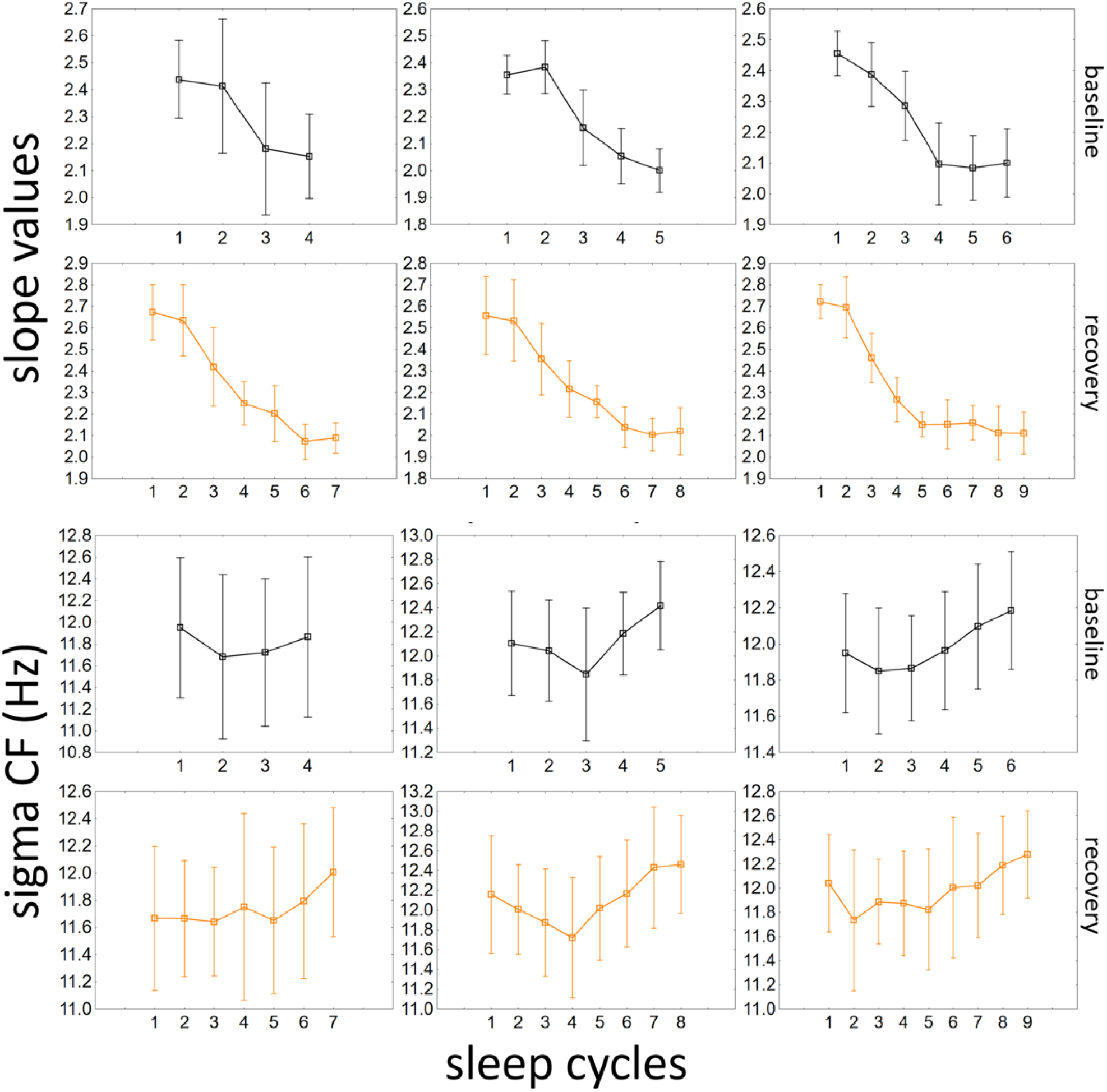
Overnight dynamics of slope values and CFs in the different cycle count-based groups. Means (open squares) and 95% confidence intervals (vertical lines) can be seen in successive sleep cycles. NREM sleep EEG spectral slopes get flatter during the night, while CF follows a decreasing trend then starts to increase toward the end of the night.

In line with our hypothesis we found no significant CF difference in the last and first cycles of sleep between BS and RS (hyp 5; *C*_*last*_: t(29)=0.12, p=0.91, *C*_*first*_: t(32)=0.89, p=0.38). Additionally, we calculated the middle time of the sleep cycle which contains the slowest CF value (NSSF). First, we tested whether NSSF reflects the subjectively measured chronotype (MCTQ MSFsc and the created chronotype groups). According to the results of One-Way ANOVA, there is a difference in BS NSSF among the different chronotype groups (F(2)=3.81, p=0.03, m_early_=2:18, SD=2:34, m_interm_=3:04, SD=2:15, m_late_=4:45, SD=1:44 hh:mm), but no difference can be captured when the RS NSSF is tested (F(2)=1.08, p=0.4, m_early_=23:07, SD=1:14, m_interm_=00:59, SD=2:08, m_late_=0:56, SD=3:17 hh:mm; Fig.3.), which results was supported by the correlational findings showing a strong positive correlation between BS NSSF and MSFsc (r=0.56, p<0.0001), but no significant correlation between RS NSSF and MSFsc (r=0.2, p=0.28). In the next step, NSSF was compared between sleep conditions to prove hypothesis 6 states that CF minima will be around the same time in both nights. However, in contrast to our assumptions (hyp 6), NSSF was significantly earlier in recovery than in BS (t(28)=5.7, p<0.0001, BS: m=03:44, SD=2:14 hh:mm; RS: m=00:40, SD=2:34 hh:mm). To examine this difference in more detail, we calculated the time of the middle of the sleep periods in BS and RS for every participant (with complete recording) and compared it with NSSF values, following the intuition that as the minimum occurs around the middle of the sleep under normal circumstances, the cause of earlier RS NSSF could be attributed to its stronger association with the sleep midpoint, rather than with the time of day. Results indicated similar NSSF and sleep period middle times in BS (as reflected by the statistically similar means and the strong positive correlation between the two metrics: *NSSF vs. Midsleep*: Z=0.97, p=0.33, Mdn_NSSF_=04:07, Mdn_Midsleep_=03:47 hh:mm; Spearman R=0.64, p<0.0001), but not in RS (significant difference: *NSSF vs. Midsleep*: t(30)=3.2, p=0.003, Mdn_NSSF_=00:11, Mdn_Midsleep_=01:46 hh:mm; nominally lower, but still significant correlation: r=0.41, p=0.02).

## 4. Discussion

The aim of the present study was to provide further evidence for the significance of NREM sleep EEG spectral parameters in estimating the fundamental processes of sleep regulation. In addition, we aimed to provide a further advancement of the idea of home sleep recordings (Korkalainen et al., 2021) by setting up an experimental intervention (sleep deprivation) taking place in the subjects’ home and not in the laboratory. Firstly, NREM sleep EEG spectral slope changed along with changes in sleep pressure (homeostatic process): it got flatter in both sleep conditions in parallel with the diminishing sleep need during the night; moreover, it got steeper in RS just as sleep pressure is increased after sleep deprivation. These findings suggest that sleep homeostasis can indeed be studied by measuring the spectral slope of the wearable headband-recorded EEG in ecologically valid, home settings. Secondly, spindle CF deceleration in the middle of the sleep period was captured in BS (with minimal CF in the middle of the night) which resulted in a U-shaped curve-like overnight dynamics of this parameter. Nonetheless, although the evolution of CF in RS was the same (CF deceleration followed by an acceleration to or above the initial CF level), the timing of minimal CF (NSSF) was neither in the middle of the sleep period, nor at the same time as that in BS.

### 4.1. Aperiodic brain activity: index of sleep homeostasis

Results of previous research works unanimously imply the brain state indexing role of spectral slope. Some of them emphasize its distinguishing function between outwardly clearly distinct brain states such as wakefulness and unconsciousness (Lendner et al., 2020). Others found proof on its fine-tuned discriminating ability within a specific brain state, like the distinction between “resting” and “working” awake brain (Höhn et al., 2024), as well as between sleep cycles (Rosenblum et al., 2022) or sleep stages (Schneider et al., 2022) in nocturnal EEG records. Our approach takes into account both these finer and larger scale findings and hypothesize that spectral slope can give an insight into the homeostatic process of sleep-wake regulation. Our former results support this claim indirectly, as we found that spectral slope is behaving similar to the gold-standard measure of sleep homeostasis the slow wave activity (SWA) during the course of regular sleep periods. Both SWA reduction and slope flattening were revealed to correlate with age, with the progression of sleep during the night, and along the fronto-posterior gradient (G. Horváth et al., 2022). In our previous reports we also discuss the advantages of slope over SWA (Bódizs et al., 2024; G. Horváth et al., 2022), which is however not the scope of the present study.

Here we provide direct evidence for slope as a homeostatic marker using a total, 35-hour long sleep deprivation protocol. In line with the prior findings, spectral slope flattened during the night in the first 4 cycles of sleep (hyp 7), and this cycle effect remained significant even when the whole sleep period was considered in the small-sample sized groups divided along the number of sleep cycles’ participants had (hyp 8). Cycle-to-cycle decrease in slope steepness proved to be significant between C2 and C3, C3 and C4 in both BS and RS, when the first 4 cycles of the sample were analysed, furthermore, between C2 and C3 in the 9-cycle group in RS. These findings are analogous to the SWA-related results of Dijk et al. (1990) reporting a similar study design with extended sleep after 36 hours of wakefulness. They found reduced SWA in consecutive sleep cycles over the first 3 cycles in BS, and over the first 5 cycles in RS after which this canonical measure of sleep intensity reached a nearly constant level. Later cycles were not found to be significantly different in terms of SWA. Note that the tendency was the same with regard to slopes in all the cycle count based groups in the present report and that no Bonferroni correction was applied in the earlier report. Comparing RS and BS, a significant condition effect and increased SWA in the first 2 cycles of RS was captured in the study of Dijk et al., (1990). When we compared slope values among conditions, we also found an overall effect of sleep deprivation (hyp 1). Slope was steeper after deprivation in the first 4 cycle of sleep, the maximum slope value indicated steeper slope in RS than in BS, however, slope minimum did not differ in the two conditions. This parallels earlier finding reporting a steady level to which SWA returns, and that early RS SWA exceeds that in BS (Aeschbach et al., 1997; Dijk et al., 1990). The maximum slope values occurred at the beginning of the sleep periods while the minima were found around the end of nights, however, in contrast to our assumptions, the steepest values were not exclusively present in the first cycle. In BS the distribution of first and second sleep cycle containing the steepest slope was almost 50-50% in the sample, however, in RS this “balance” shifted towards the first cycle. Indeed, the pattern of overnight flattening of the sleep EEG spectral slopes reported herein strongly resembles the decline of 0.3–0.9 Hz band-limited power across NREM periods 1–4, both following a convex curvature (Campbell et al., 2006). Accordingly, the lack a significant difference between <1 Hz NREM sleep EEG power of the first and the second sleep cycle was reported repeatedly (Achermann & Borbély, 1997; Borbély et al., 1981). As we intentionally avoided fitting on the spectral peak region of the slow (<1 Hz) oscillation, looking for the spectral slopes between 2 and 48 Hz, the above detailed resemblance implies an apparent contradiction. However, several lines of evidence suggest that the cortical bistability characterized by up-down state alternation at roughly 1 Hz frequency results in fractal spectrum in a wide frequency range exceeding significantly the <1 Hz power (Baranauskas et al., 2012; Bódizs et al., 2024; Milstein et al., 2009). As a consequence, this finding might indicate the preferential contribution of cortical bistability to the emergence of NREM sleep EEG spectral slope or fractal spectra. Another possible explanation for the lack of a significant difference between the first and the second NREM period in terms of EEG spectral slopes is the “first use effect” of the headband EEG device in BS, which could disturb the very beginning of sleep resulting in slightly flatter slopes in the first compared to the second sleep cycle.

### 4.2. Oscillatory frequency in the spindle range

Oscillatory frequency of sleep spindles was suggested recently as an EEG-biomarker for circadian rhythm by the Fractal and Oscillatory Adjustment Model (Bódizs et al., 2024) of sleep regulation. This suggestion was based on several findings supporting time-of-day modulation of spindles (Dijk et al., 1997), its association with validated circadian markers (Knoblauch et al., 2005; Wei et al., 1999), and its change during the lifespan (Bódizs et al., 2022; Purcell et al., 2017; Wei et al., 1999) in parallel with the age-related changes in circadian biology (Duffy et al., 2015; Roenneberg et al., 2004).

When the overnight dynamics in CF of the two nights were tested separately for complete recordings, the cycle effect was significant in most groups (hyp 10), but not when the first 4 cycle of the entire sample were analysed (hyp 9). This is logical considering the expected behaviour of CF mid-sleep deceleration and that more than half of the participants had more than 4 sleep cycles already in BS (Mdn_BS_=5 cycle). To capture mid-sleep deceleration of CF, a comparison of the first and last cycle with the middle cycle was conducted. CF was found to be significantly faster in the last cycles of sleep than in the middle, but there was only a tendency regarding first-to-middle cycle deceleration in both sleep conditions. Furthermore, when we compared first with last cycles separately in the two conditions, we found that last CF values were significantly faster than first ones. The pattern of CF development throughout the night indeed following a U-shape, and the deceleration in the first half of the night is a consequent finding in earlier reports (Bódizs et al., 2022; G. Horváth et al., 2022, also see the development in dominant frequency changes during the night in in Aeschbach et al., 1997), but not always a statistically significant change. It is possible that spindle frequency peaks somewhere in the first half of the biological day, which is supported by findings showing that spindle frequency is higher in daytime sleep compared to that occurring during night time (Knoblauch et al., 2003, 2005; Rosinvil et al., 2015; Wei et al., 1999).

However, the long-term sleep deprivation might also have an impact on the partially inconsistent findings regarding RS in the present study. Although CF correspondence in the last and first cycles of the different sleep conditions was fulfilled (hyp 5), CF development in the first 4 cycles was not different between the sleep conditions (hyp 4). Furthermore, there were large discrepancies regarding the assumed phase-indicator, NSSF. On the one hand, we hypothesized that overnight CF trajectory will differ between conditions, as the bedtime was earlier, and as sleep period was longer for all participants in RS. Besides, we assumed that NSSF will be around the same time in the two nights (hyp 6) suggesting a time-of-day dependency of this variable. However, the tendency of CF deceleration started already in the second cycle of sleep in RS (such as in BS), and NSSF occurred significantly earlier in RS than in BS. Furthermore, although BS NSSF was reflected in subjective chronotype, and occurred around the middle of the night, this was not true for NSSF in RS. One possible explanation can be the severity of the intervention. 35 hours of wakefulness challenges the homeostatic process to such extent that the regulation of circadian pacemaker may be overshadowed. Indeed, in their recent literature review, Franken and Dijk (2024) concluded that the two processes are much more entangled as compared to the basic assumption expressed in the original two process model. Furthermore, data on biomarkers of the circadian rhythm seems to support the above entanglement. The phase-advancing effect of morning bright light measured in terms of Dim Light Melatonin Onset (DLMO) indices was found to be reduced after acute, short-time sleep deprivation (Burgess, 2010), whereas a delayed acrophase of melatonin and suppressed BMAL1 gene expression after one night sleep deprivation was found (Ackermann et al., 2013). The circadian phase shifting effect of darkness was also reported (Buxton et al., 2000; Santhi et al., 2005). As in the present study a lights-off instruction was given to the participants at bedtime, early darkness in RS could advance the circadian phase of the participants, which could result in earlier NSSF. Besides, there is a study that measured electrical activity of the suprachiasmatic nucleus (SCN; the pacemaker of the circadian timing system) along with EEG and found that neuronal activity of SCN depends on the vigilance state of the animal, for instance, NREM sleep is lowering it (Deboer et al., 2003). This finding supports the idea that earlier sleep window can modulate CF development even if it is primarily related to the circadian process (like SCN-activity). Given that CF-related variables in BS mostly behaved as expected, such as U-shape dynamics (mid-sleep deceleration), or the associations between subjectively measured chronotype and BS NSSF, we think that after all the advanced sleep protocol was rather a limitation of the time-of-day analyses in this study. Thus, further studies with the direct modulation of the circadian cycle, with the follow up of spindle frequency development in the whole 24-hour period are needed.

### 4.3. Limitations and overall conclusions

Overall, our findings corroborate the idea of Fractal and Oscillatory Adjustment Model, thus support the use of spectral slope as a homeostatic marker, and highly recommend further research on spindle frequency as a circadian phase indicator. Among the limitations we can mention the lack of adaptation night, and the advanced sleep protocol which latter contradicted our expectations regarding the time-of-day analyses as outlined above. However, despite further constraints such as the availability of frontolateral EEG recordings only, small group sizes, or limited age-range, results are clear regarding spectral slope as an indicator of sleep homeostasis, and promising about CF.

## 4.4 Acknowledgments

This research has been implemented with the support provided by the Ministry of Innovation and Technology (TKP2021-EGA-25 and TKP2021-NKTA-47) and supported by the ÚNKP-23-3-II New National Excellence Program of the Ministry for Cultural and Innovation from the source of the National Research, Development and Innovation Fund.

